# Reversible supramolecular assembly of the anti-microbial peptide plectasin into helical non-amyloid fibrils

**DOI:** 10.1101/2021.09.06.458672

**Authors:** Christin Pohl, Gregory Effantin, Eaazhisai Kandiah, Sebastian Meier, Guanghong Zeng, Werner Streicher, Günther H.J. Peters, Guy Schoehn, Christoph Mueller-Dieckmann, Allan Noergaard, Pernille Harris

## Abstract

Self-assembly and fibril formation play important roles in protein behavior. Amyloid fibrils formation is well-studied due to its role in neurodegenerative diseases and characterized by refolding of the protein into predominant β-sheet form. However, much less is known about the assembly of proteins into other types of supramolecular structures. Using cryo-electron microscopy at a resolution of 1.97 Å, we show that a triple-mutant of the anti-microbial peptide plectasin assembles reversibly into helical non-amyloid fibrils. Plectasin contains a cysteine-stabilized α-helix-β-sheets structure, which remains intact upon fibril formation. Two fibrils form a right-handed superstructure with each fibril consisting of double helical, left-handed structures. The fibril formation is reversible and follows sigmoidal kinetics with a pH-dependent equilibrium between soluble monomer and protein fibril. The anti-microbial activity does not appear compromised by fibril formation. This is the first high-resolution structure of this type of α/β protein fibrils.

## INTRODUCTION

Self-assembly is a hallmark of biomacromolecules. For protein polymers and fibrils, assembly can be highly ordered and can serve different purposes. Protein fibrils have been found to serve as structural elements in long-term memory in microorganisms^1,2^. In mammalian cells, they were also found to template Melanin polymerization as a protection against cytotoxic damage^3^, to constitute part of peptide storage in the endocrine system^4^ and to be involved in signal transduction^5^. Many polymeric protein assemblies found in nature are amyloid fibrils that are characterized by a distinct cross-β sheet motif.^6^ Amyloids are best known for their role in neurodegenerative diseases like Alzheimer’s disease^7^ or prion related diseases like Creutzfeldt-Jakob disease and Bovine Spongiform Encephalopathy^8^. Recent studies have shown a strong link between anti-microbial peptides (AMPs) and amyloid proteins. Amyloid-β, which is known to form amyloid fibrils associated to Alzheimer’s disease, has been found to have anti-microbial and antifungal activity linked to the innate immune system^9–11^. Several other amyloids or their fragments, including α-synuclein^12^, islet amyloid polypeptide (IAPP)^13–15^, tau protein^16^, human prion protein^17^ and endostatin^18^ have been reported to show anti-microbial activity. Furthermore, many AMPs, including, lysozymes^19^, protegrin-1^20^, HAL-2^21^, uperin 3.5^22^, dermaseptin S9^23^, Cn-AMP2^24^ and longipin^25^ have been found to form amyloid or amyloid-like structures. In a direct comparison, similar mechanisms of membrane disruption have been found for amyloids and AMPs^14,26^. Dysregulation of amyloid proteins leading to amyloid fibrils are mainly associated with diseases or other negative effects^27,28^. Moreover, dysregulation of AMPs has been associated with chronic inflammation and autoimmunity reactions^29–31^.

Amyloid fibrils have been studied intensively over the last century. The study on fibril formation and self-assembly by non-amyloid proteins has remained much more limited due to their lower natural abundance compared to amyloid fibrils. The study of anti-microbial non-amyloid polymers has mainly focused on their synthetic design to overcome challenges in immunogenicity, off-target effects and serum instability and ensure their controlled release^32–34^. Furthermore, the self-assembly of natural AMPs has been mainly claimed to be of amyloid nature^21,22,35^. However, in recent years several high-resolution structures revealed assemblies that differed from the typical cross-β assembly in amyloid fibrils^36–38^.Recent developments in cryo-electron microscopy (cryo-EM) have proven valuable in the structure determination of insoluble protein fibrils. A variety of different protein fibril structures have been revealed, including cross α-helical amyloid-like fibrils^36,37^ and functional α-helical assemblies of the anti-microbial LL-37^38^.

Plectasin is an endogenous peptide with a molecular weight of 4.4 kDa that show antibiotic behavior against Gram-positive bacteria^39^ by binding lipid II and preventing its incorporation into the bacterial cell wall^40^. Plectasin consists of 40 amino acids and shows well-defined secondary structural elements in form of an α-helix (M13-S21) and an antiparallel β-sheet (G28-A31; V36-C39)^39,41^. The structure of plectasin is stabilized by three disulfide bridges (C4-C30; C15-C37; C19-C39), which is typical for this type of defensins and leads to a very high conformational stability^42^. Its antibiotic behavior makes plectasin a protein drug candidate in the efforts against increasing bacterial resistance, as plectasin showed favorable characteristics such as low toxicity, high serum stability and long *in vivo* half-life^39,40^ in addition to its anti-microbial activity.

Here, we describe the non-amyloid fibril formation of the plectasin variant D9S Q14K V36L, referred to as PPI42 using cryo-EM and biophysical techniques. Previously, we could show that the introduced mutations, which were originally implemented to increase anti-microbial activity, were predicted by MD simulations to only have a minor effect on the overall structure of plectasin^42^ (Figure S1). We report the structural details of the fibrils, the pH-dependence and kinetics of fibrils formation. PPI42 displays a novel fibril structure comprised of the native-like protein forming a left-handed double helical structure. We show that two fibrils subsequently assemble into a right-handed superstructure. To the best of our knowledge, this is the first example of an atomic-level characterization of fibrils in equilibrium with a sparsely soluble monomer formed by an α/β defensin.

## RESULTS

### Plectasin variant PPI42 forms a gel with regular fibril superstructure

Plectasin variant PPI42 formed a gel at peptide concentrations of approx. 5 mg/mL and higher, during dialysis at pH 5 and above, whereas the plectasin wildtype remained in solution under the same conditions. Gel formation occurred irrespective of the chemical nature of the buffers tested in this study (histidine, tris, acetate, citrate, or no buffer). When dialyzed against acidic pH, the gel formation proved reversible, and the gel dissolved into a liquid. In our previous study, we found a pH dependent loss in nuclear magnetic resonance (NMR) signal intensity of PPI42 at constant protein concentration and an exponential increase of apparent molecular mass and radius of gyration *R_g_* with increasing protein concentration in small-angle X-ray scattering (SAXS) measurements^42^, indicating that large protein clusters formed and resulted in gel formation.

Using atomic force microscopy (AFM), the nanoscale structure of the gel was investigated by PeakForce Quantitative Nanomechanical Mapping (QNM™). This mode made simultaneous mapping of topography, and elasticity of the sample possible. The topography of the gel formed by PPI42 at pH 5 was investigated at peptide concentrations of approx. 20 mg/mL (4.5 mM) (Figure 1A). Long pearl necklace-like fibrils could be observed. Different buffer systems were tested, but no significant differences in the topography were observed (Figure S2). The height of the fibrils in different buffers was measured to 3.83±0.30 nm with a periodicity of 29 nm. All samples showed similar morphology and comparable elasticity of 0.7-1.0 GPa, indicating that the fibrils formed by PPI42 were independent of the buffer and possibly suitable for high resolution structural studies by electron microscopy (EM).

**Figure 1:**
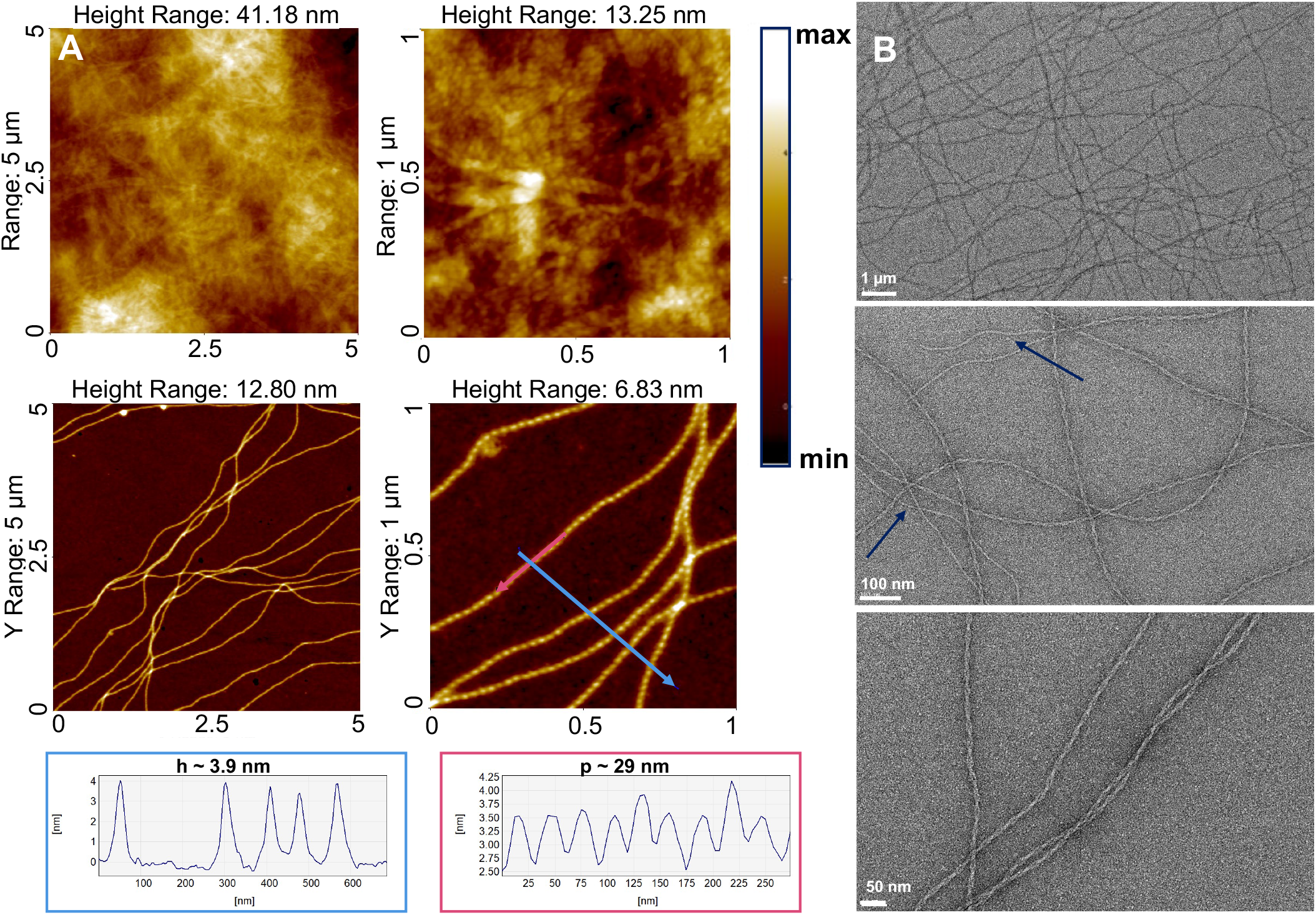
A: AFM height images of various ranges and line profiles measured along the shown arrows of PPI42 dialyzed at pH 5. The line profile in the blue box corresponds to measurement along the blue arrow, the line profile in the red box corresponds to measurement along the red arrow. B: Images of the fibrils formed by PPI42 imaged using negative stain EM. PPI42 fibrils formed a right-handed helical superstructure consisting of two single strands. Frayed fibril termini and single fibrils are indicated by arrows.

Negative stain EM was used to gain more insights into the structure of the fibrils formed by PPI42. The fibrils were formed at pH 5.5 and the total peptide concentration after dialysis was ~20 mg/mL (4.5 mM). EM measurements revealed that the pearl necklace-like structure derived from AFM actually consisted of two single fibrils winding around each other to form a right-handed helical superstructure (Figure 1B). We observed frayed fibril termini of and a small number of single fibrils, suggesting that the single fibrils of the PPI42 formed first and then assembled into the helical superstructure. The network built by PPI42 fibrils showed similar features in AFM and negative stain EM (Figure 1). The periodicity of 29 nm along the fibril obtained by negative stain EM (Figure S3) was very similar to the periodicity measured with AFM.

### CryoEM Structure of the Supramolecular Assembly

We were able to determine the structure of the plectasin fibril at a resolution of 1.97 Å using cryo-EM, which was one of the highest resolutions hitherto observed for a peptide fibril in cryo-EM and allowed an unambiguous structure determination (Figure 2, Figure S4). The high-resolution structure confirmed the right-handed helical superstructure consisting of two peptide fibrils observed in negative stain EM. Within each fibril, PPI42 assembled into a double stranded left-handed helix with an axial rise of 25.2 Å and a helical twist of 15.75°. The asymmetric unit was composed of seven monomers arranged in a near-helical way with an average axial rise of 3.75 ± 0.15° and an average twist of 156.48 ± 2.25° (Figure S5) with variations resulting from interactions of the two fibrils leading to a curvature. The axial rise appeared, when measured in X-ray fiber diffraction, as a sharp diffraction ring at ~ 4 Å (Figure S6). Weaker diffraction rings were observed at 6.2, 8.0 and 9.0 Å, which corresponded most likely to different distances between the monomers. The overall structure of PPI42 within the fibril proved to be similar to the plectasin wildtype and showed only minor differences to our simulated structure of PPI42 in solution (all-atom RMSD: 1.059 Å)^42^. The N-terminal loop and the loop connecting the antiparallel β-sheet were oriented towards the fibril center while the C-terminus and the loop between α-helix and β-sheet were facing outwards. PPI42 fibril was predominantly stabilized within the same strand both by polar and hydrophobic interactions (Figure 3). Each PPI42 monomer *i* interacted with its two directly flanking monomers *i*−1 and *i*+1 in the same strand (Figure 2). The interactions between the strands occurred between a PPI42 monomer *i* and three monomers in the other strand *n* and its adjacent monomers *n*−1 and *n*+1. The negatively charged sidechains of E10 and D11 played an essential role in the arrangement of plectasin to the fibril by forming a network of salt bridges to different lysines (Figure 3B). Residue (D11)_*i*_ formed a salt bridge within the same strand to the (K14)_*i*−1_ sidechain. Additionally, (D11)_*i*_ formed a salt bridge connecting the two different strands to (K32)_*n*_. Residue (E10)_*i*_ also connected the two strands by forming a salt bridge to (K32)_*n*+1_ and (K38)_*n*+1_. On the opposite site (N5)_*i*_ formed a hydrogen bond to (G6)_*i*+1_ backbone within the same strand. At the center of the fibril (W8)_*i*_ formed a hydrogen bond to (G33)_*n*_ backbone in the neighboring strand. In addition to these polar interactions, the PPI42 fibril was stabilized by a hydrophobic core in the fibril center of each double strand and hydrophobic interfaces between the monomers within each strand (Figure 3A).The hydrophobic center consisted of residues P7, W8 and F35 (Figure 3A, Figure S4). CH/*π*-interaction between P7 and W8 additionally stabilized the fibril center (Figure 3C). Hydrophobic interactions on the fibril surface consisted of (F2)_*i*_ and (H18)_*i*_, which were stabilized by aromatic π-stacking, clustering with M13 and H16 on molecule *i*+1 (Figure 3D), resulting in a series of interfaces that were located in a ring-like manner (Figure 3A). The assembly of the two protein fibrils into the superstructure is favored by the interaction of the loop between α-helix and β-sheet of two consecutive monomers *i* and *i*−1 in the same strand with two symmetric monomers *m* and *m*+1 in the other fibril. The (K23)_*i*_ sidechain and backbone and (S21)_*i*_ backbone formed three hydrogen bonds with (G24)_*m*_ and (K23)_*m*_ backbone which stabilized the superstructure (Figure 3E).

**Figure 2:**
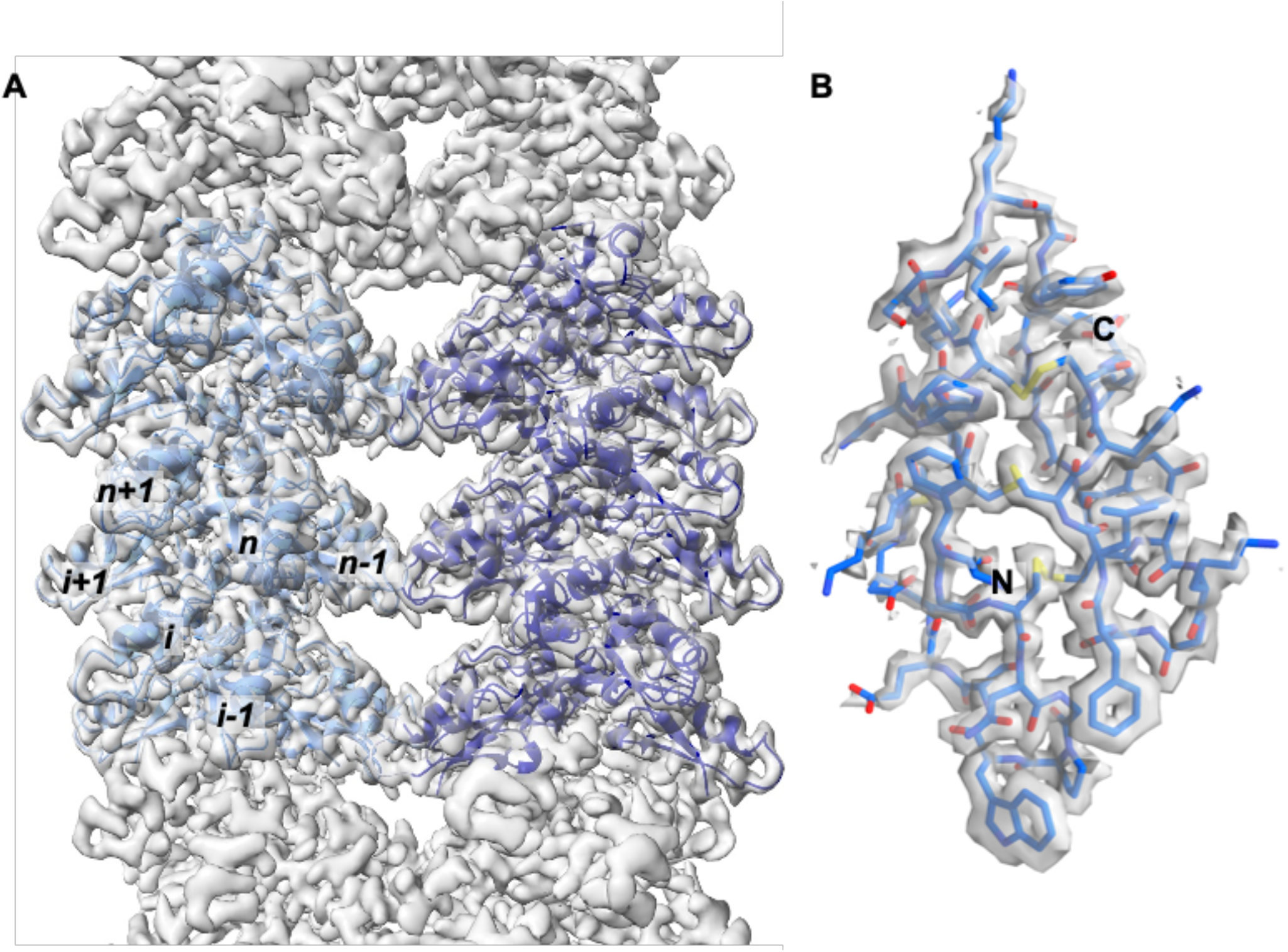
High-resolution cryo-EM structure at 1.97 Å resolution of the protein fibril formed by PPI42. A: Cryo-EM map including modelled monomers of the right-handed superstructure consisting of two fibrils with left-handed double helical symmetry. B: Atomic model of PPI42 built into the electron density.

**Figure 3:**
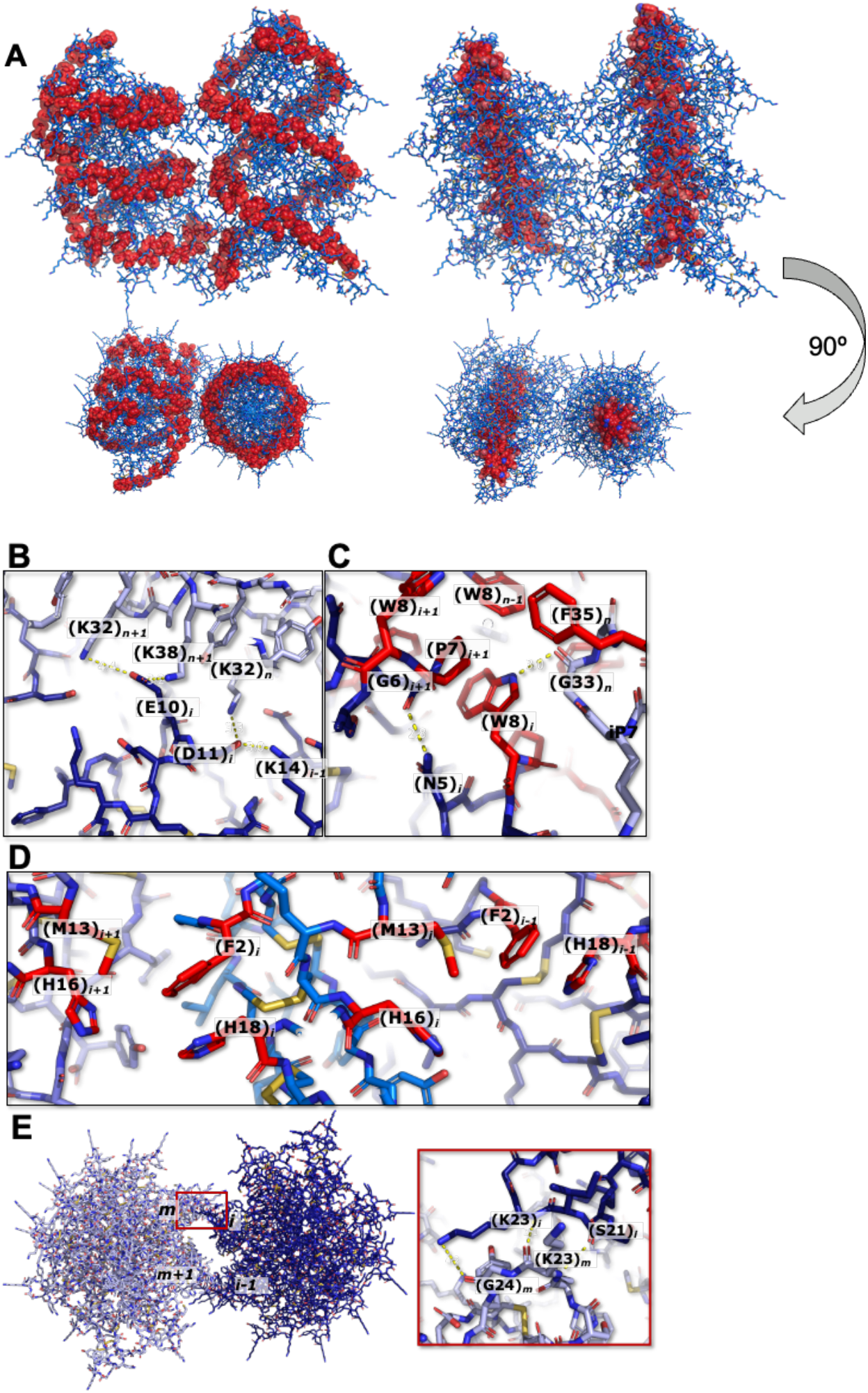
Stabilizing protein-protein interactions within superstructure of PPI42 fibrils. A: Stabilizing hydrophobic interactions within PPI42 fibrils forming an outer ring, consisting of F2, M13, H16 and H18 and a hydrophobic center, consisting of P7, W8 and F25. Hydrophobic amino acids forming the clusters are shown as red spheres. View along the transverse axis of the fibril and view along the longitudinal axis of the fibril is shown (90º rotation). B: Polar interactions (yellow lines) of acidic amino acids (E10)_*i*_ and (D11)_*i*_ within one strand with (K14)*i*−1 of the adjacent monomer and between the strands with (K32)_*n*_ and (K32)_*n*+1_ and (K38)_*n*+1_. C: Polar (yellow lines) and hydrophobic (red amino acids) interactions in the fibril center. D: Hydrophobic interactions (red amino acids) within one strand forming the outer hydrophobic ring of the PPI42 fibrils. E: Polar interactions (yellow lines) between the two fibrils forming the right-handed superstructure with view along the longitudinal axis of the fibril.

The high-resolution structure of PPI42 forming a fibril was determined using the superstructure. In our negative stain EM measurement, we observed a small portion of fibrils existing as single strands. In order to investigate whether there are structural rearrangements between fibril and the fibril superstructure, we analyzed the 3.3 Å resolution structure of the single strand obtained by cryo-EM (Figure S4). PPI42 shows a very similar structure and assembly in the single strand compared to the superstructure (all-atom RMSD: 0.727 Å) (Figure S5). PPI42 showed a strict helical symmetry in the single fibril with an axial rise of 3.76 Å and a helical twist of 156.5 Å, which is almost identical to the helical arrangement in the superstructure.

To elucidate the structural features that allow PPI42 to assemble into fibrils, while the plectasin wildtype did not fibrillate, we determined the crystal structure of the plectasin wildtype at a resolution of 1.1 Å (Figure S5). Our plectasin wildtype structure showed a high similarity to the published crystal structure (PDBID: 3E7U^43^) with only minor differences in the N-terminal loop (overall 0.316 Å RMSD). When we compared our crystal structure of the wildtype plectasin to the structure of PPI42 within the fibril, we observe a significant structural difference in the N-terminal loop region between amino acids 9 and 14 (Figure 4A). Both positions are mutated in PPI42 (D9S; Q14K). Therefore, we analyzed the coordination within this region in detail. We observed a network of polar interactions in the plectasin wildtype with D9 forming a salt bridge to N5. When mutated to S9 in PPI42, the sidechain orientation changes and the sidechain forms a hydrogen bond to G6 backbone, which disrupted the network stabilizing this part of the N-terminal loop (Figure 4B). Additionally, the N5 sidechain was not internally coordinated, making the formation of polar interactions with neighboring proteins possible. Notably, the sidechains of E10, D11 and D12, formed intermolecular interfaces both in the protein crystal of the wildtype and in the fibril of PPI42. Their orientation differed, however, significantly between wildtype and PPI42 (Figure 4C). This structural difference may contribute to the different arrangement with the mutant forming the fibril and the wildtype forming the crystal. The Q14K mutation additionally led to a different coordination of H18 (Figure 4D), which played an important role in forming the hydrophobic outer ring in the fibrils. In the plectasin wildtype, H18 was coordinated in two different ways (50/50 distribution) stabilized by polar interaction with Q14 and *π*-stacking with F2, while in PPI42 only the *π*-stacking with F2 occurred due to the mutation Q14K.

**Figure 4:**
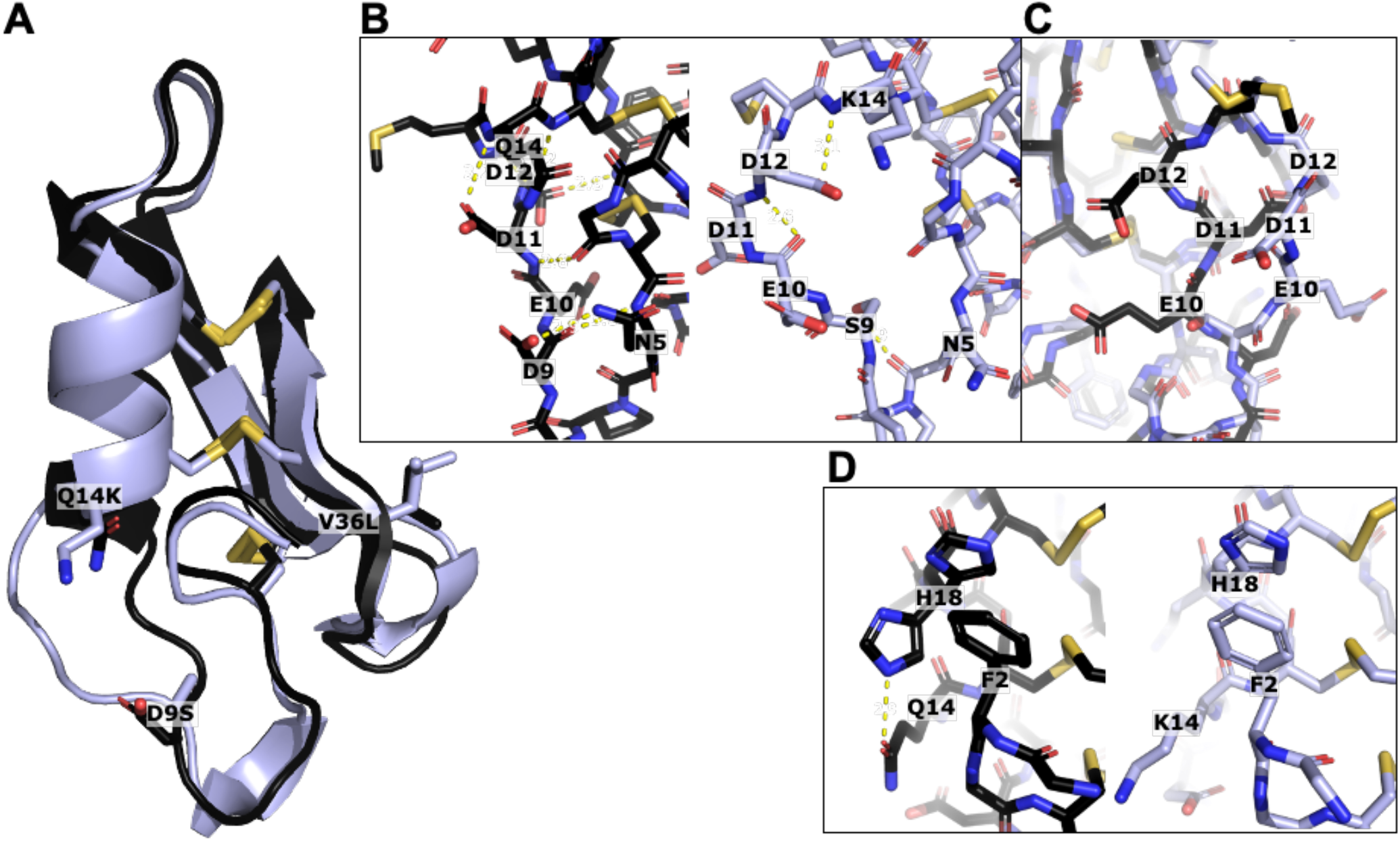
Comparison of the plectasin wildtype crystal structure (black) and PPI42 fibril structure (light blue). A: Overall comparison of the two structures shown as cartoon. The three mutated amino acids are highlighted as sticks. B: Polar coordination of the N-terminal loop in the plectasin wildtype (left) and PPI42 (right). C: Direct comparison between side chain orientation of the acidic amino acids E10, D11 and D12 in wildtype (black) and PPI42 (light blue). D: Differences in the coordination of H18 due to Q14K mutation leading to two distinct orientations of H18 in the wildtype (left) compared to PPI42 (right).

### Optical studies of fibrillation onset and mechanism

As Rayleigh scattering is proportional to the sixth power of the particle radius, it readily captures small fractions of large oligomers. Hence, dynamic light scattering (DLS) was used to determine the onset and concentration dependence of fibril formation. The pH onset was measured at low peptide concentration of 2 mg/mL (450 μM), where PPI42 did not form a gel. Based on the results, three size ranges for the DLS measurements were set, 0.1-10 nm to capture the monomeric fraction, 10-100 nm to capture medium sized oligomers and 100-1000 nm to capture the fibrils. At pH 5 and a peptide concentration of 2 mg/mL, PPI42 showed no significant scattering from oligomers. From the cumulant analysis method of the autocorrelation function a hydrodynamic radius *R_h_* of 1.53 nm was determined, which is close to the calculated monomeric *R_h_* of 1.37^44^ nm based of the monomer in the fibril, confirming that this fraction represented mainly monomeric species. Above pH 6, we observed a small fraction (<2.5% mass) of medium sized oligomers and protein fibrils (Figure 5A). Therefore, we measured the concentration dependency of the formation of these large oligomers or aggregates at pH 6.5. We observed a concentration dependent increase of the fraction of fibrils. The fraction of medium sized oligomers did not increase with concentration indicating a direct addition of monomers to the fibril. Analysis of the mass percentage of each fraction showed a similar trend (Figure S6). This measurement indicated a pH and concentration-dependent equilibrium between monomer and peptide fibril. The strong scattering influence of large polymers in DLS measurements implied that small variations in quantities of fibrils or other large particles can result in issues with exact quantification and reproducibility, which led to high standard deviations in our measurements.

**Figure 5:**
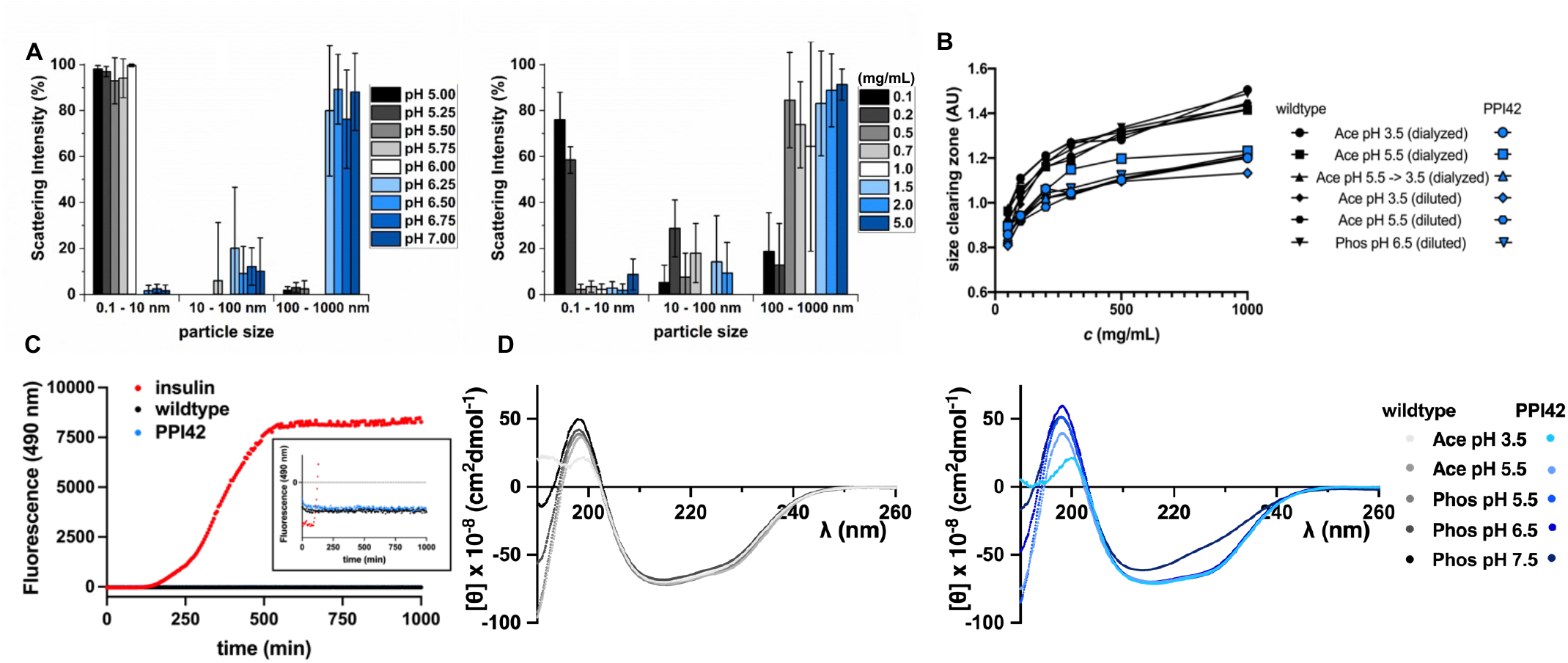
Molecular characteristics of the protein fibrils formed by PPI42. A: Dependence of fibril formation on pH (c = 2 mg/mL, left) and protein concentration (pH 6.5, right) determined by DLS. The displayed data are mean values ± S.D. for 3 replicates. B: Anti-microbial activity of the plectasin wildtype and PPI42 measured by the size of the clearing zones in the semi-quantitative radial diffusion assay. The anti-microbial activity was measured as a function of peptide concentration in different conditions relevant for this study. C: Thioflavin T (ThT) fluorescence measurement over time on plectasin wildtype, PPI42 and insulin, which was used as a positive control (*c*=2 mg/mL, *T*=40°C). Insert shows ThT fluorescence monitored over time of wildtype and PPI42 in detail. D: Far-UV CD spectra of the plectasin PPI42 (blue, left) and wildtype (right, black) and as a function of pH (c=0.132 mg/mL (30 μM)).

### Plectasin function and structure are weakly affected by fibrillation

In order to investigate whether the fibril formation has an impact on the anti-microbial activity of PPI42 we performed a semi-quantitative radial diffusion assay (RDA). The anti-microbial activity of the plectasin wildtype and PPI42 was tested against *Staphylococcus carnosus* in different conditions as a function of protein concentration (Figure 5B, Figure S6). PPI42 showed generally smaller clearing zones than the plectasin wildtype. However, no large differences in the size of the clearing zones could be observed between the tested conditions, despite PPI42 formed a gel at pH 5.5 and higher. Additionally, we tested a sample that was first dialyzed at pH 5.5, where gel formation occurred for PPI42, and subsequently dialyzed into pH 3.5. No differences in their anti-microbial activity were observed, indicating that fibril formation did not hinder the anti-microbial activity. It remains, unclear whether PPI42 anti-microbial activity is only attributed to monomeric peptides or to peptides in their fibrillose state. Nevertheless, notwithstanding, supramolecular fibrils of plectasin may have the potential to provide a slow release of the monomer, which is known to be bioactive.

We subsequently used circular dichroism (CD) spectroscopy to investigate whether PPI42 secondary structure was compromised upon fibril formation, and if the observed structural differences of PPI42 compared to the wildtype were only present in the fibril form. The far-UV CD spectra of wildtype and PPI42 were investigated as a function of pH (peptide concentration of *c*=0.132 mg/mL (30 μM)) (Figure 5D). The spectra of PPI42 and plectasin wildtype looked very similar except for pH 7.5, where the CD spectrum of PPI42 showed a clear change. However, the CD spectrum still resembled the presence of α-helix and antiparallel β-strands, indicating that the overall secondary structure remained intact. The observed change in the spectrum resulted most likely from a change in the N-terminal loop region as observed in cryo-EM, which affected the secondary structure. The observed change in CD-spectrum was small in comparison to structural changes observed when amyloid fibrils are formed^47^.

Thioflavin T (ThT) is widely used to follow amyloid fibril formation via fluorescence measurements^45^ and has shown potential to follow other types of protein fibrils^36,37^. We investigated therefore whether the fibril formation could be followed by ThT fluorescence measurements despite the native-like protein structure of PPI42 (Figure 5E). Insulin, which is known to form amyloid fibrils upon shaking at 40°C^46^, was used as a positive control. Insulin, plectasin wildtype and PPI42 were investigated at peptide concentrations of 2 mg/mL. In contrast to insulin, neither the wildtype sample, nor PPI42 showed increased fluorescence. Since we observed protein fibrillation to be pH responsive, we tested different buffer conditions, but no fluorescence increase could be observed, opposite to an equilibrated insulin sample (Table S1). Additionally, different temperatures and forms of mechanical stress were tested, but none led to an increase of fluorescence. These findings were consistent with the fact that PPI42 forms non-amyloid fibrils.

### In situ NMR spectroscopy of factors affecting fibrillation

We utilized real-time NMR spectroscopy to follow the kinetics of PPI42 fibrillation *in situ* at various pH values and at a peptide concentration of 1.1 mg/mL (250 μM and 50 mM phosphate buffer). Due to their large size and long rotational correlation time, fibrils of PPI42 are not visible in liquid state NMR, and the time series therefore followed the fraction of monomer over time (Figure 6A). The kinetics of protein fibrillation were followed based on integration of the methyl groups of I22 and L36 (Figure S7). The kinetic progress of fibrillation notably showed sigmoidal behavior, strongly indicative of the need for an initial nucleation step to facilitate fibril growth. Additionally, both the fibrillation kinetics and the fraction of monomer in solution at equilibrium were pH dependent. The pH-responsive fibrillation kinetics showed a transition between pH 6.0 and 7.0, indicative of the titration of a histidine residue playing a role in fibrillation. The titration behavior of chemical shifts for both protons in the imidazole rings of H16 and H18 was therefore determined by 2D ^1^H-^1^H TOCSY NMR spectra (Figure 6B). The *pK*_a_ for H16 was around 4.4, while the *pK*_a_ for H18 was around 6.4, which corresponded to the pH where fibril formation was observed. H18 and H16 are involved in the hydrophobic interactions on the monomeric interfaces forming of the outer hydrophobic ring, which might explain the pH responsive fibril formation of PPI42 upon sidechain deprotonation. S21 is involved in the interactions between the two fibrils and showed a similar pH response as H18, indicating that the helix cap is structurally affected when fibrils are formed.

**Figure 6:**
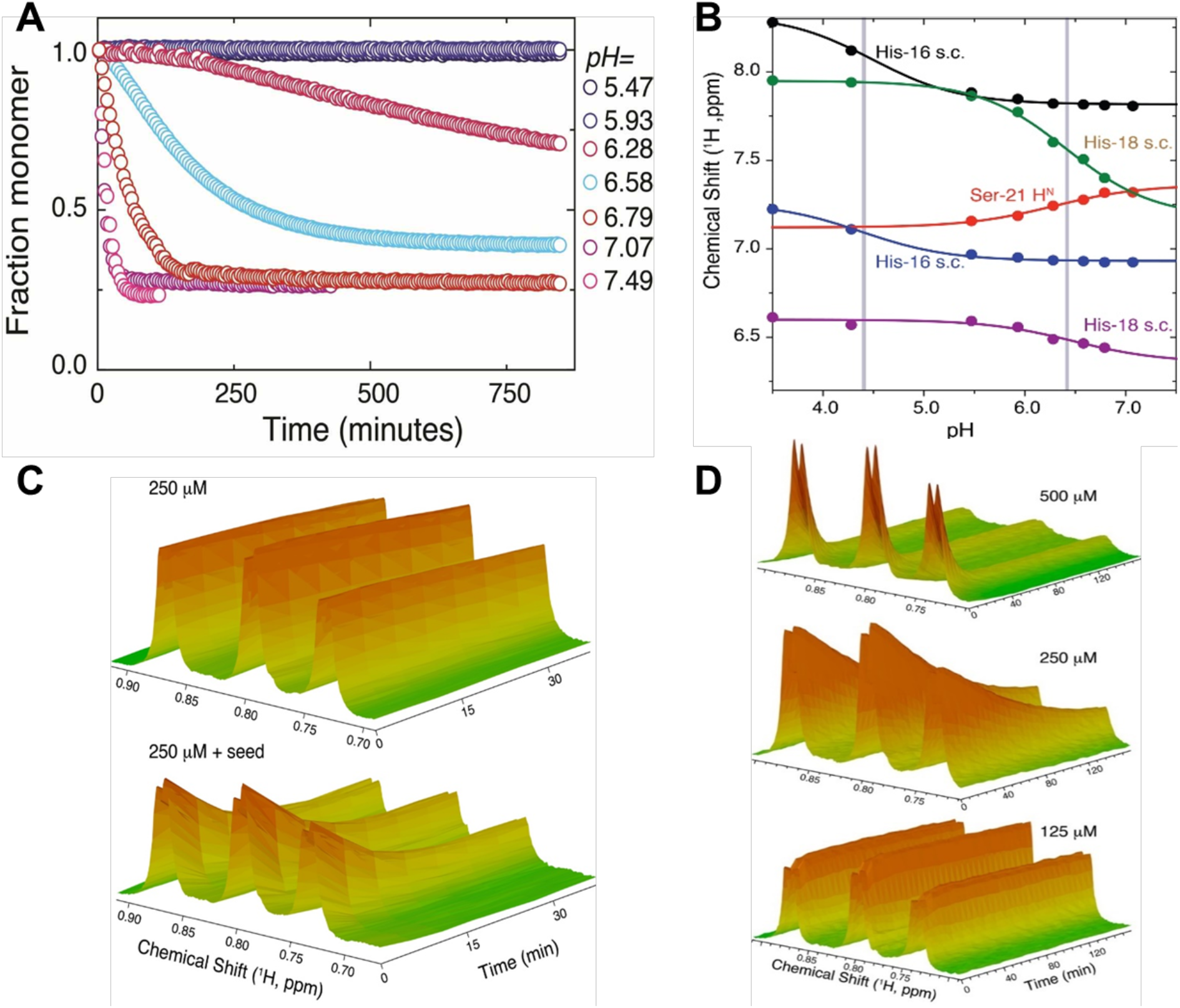
Fibrillation kinetics of PPI42 measured with NMR. A: The fraction of monomer was determined by integration of ^1^H signals for the I22 and L36 methyl groups relative to time 0. B: Titration curves of PPI42 H16, H18 (H^2^ and H^4^ in imidazole ring) and S21 H^N^ determined by identifying the signals with 2D ^1^H-^1^H TOCSY. The grey bars indicate the *pK_a_* of the titratable side chains. All measurements were conducted in 50 mM phosphate buffer. C: Fibrillation kinetics with and without seeding with 5 μl of an equilibrated sample at pH 6.75. D: Concentration dependence of fibrillation kinetics at pH 6.75

Subsequently, fibrillation kinetics were investigated with regards to dependence on seeding and monomer concentration. To a sample of PPI42 freshly diluted into pH 6.75, 5 μL of a previously equilibrated sample in the same condition was added for seeding, which led to a vastly accelerated fibrillation (Figure 6C). This observation supported that a nucleation step occurs in fibrillation. Fibrillation kinetics as followed by *in situ* NMR at varying peptide concentrations at pH 6.75 finally showed that the supramolecular reaction had a strong dependence on protein concentration (Figure 6D).

## DISCUSSION

Here, we determined the structure and kinetics in the pH-responsive self-assembly of PPI42 into non-amyloid protein fibrils. With a combination of AFM and negative stain EM we characterized the network and surface morphology of the fibrils. The fibril network observed was consistent with negative stain EM and AFM measurements and the dimensions and elasticity were similar in all different buffers, showing that the different buffer systems, ionic strengths (Table S2) or treatments did not have any significant effect on the fibrils, which is considered advantageous for medical applications. The high-resolution structure of the peptide fibrils formed by PPI42 solved at a resolution of 1.97 Å revealed that native-like monomers assemble into a left-handed double helical structure. The structure differs from the recently reported high-resolution structure of the anti-microbial LL-37^38^ and is the first example of this type of fibril. The structure was stabilized both by polar as well as hydrophobic inter-subunit interactions. An outer hydrophobic ring may explain the structure of PPI42 fibrils. Polar interparticle interactions within the acidic amino acid patch (D9-D12) were played a significant role in PPI42 fibrils and crystal interactions of plectasin wildtype. Within this patch, we observed significant conformational differences between wildtype and PPI42 structure, whereas the remaining part of the peptide proved structurally very similar. This might explain the different types of self-assembly of wildtype and PPI42. A comparison between the cryo-EM structure of the superstructure and of the single fibrils, determined at 3.3 Å resolution revealed a high structural identity, consistent with the formation of single fibrils prior to the assembly into a curved superstructure. The elasticity measured for PPI42 fibrils (0.7 and 1.1 GPa) was lower than typically observed for amyloid fibrils (2-4 GPa)^48,49^, but is comparable to non-amyloid fibrils formed by ovalbumin^50^.

Supramolecular association of PPI42 into fibrils was pH responsive, which has been reported for other self-assembling peptides^51,52^. The equilibrium distribution of PPI42 between monomers and fibrils was dependent on pH and peptide concentration. Increasing either of these two parameters shifted the equilibrium towards fibrils. Hence, highly concentrated sample could self-assemble at a lower pH than lower concentrated sample, explaining different pH onsets observed in this study. The small fraction of soluble, medium sized oligomer did not show a clear trend with increasing pH or concentration. These results suggest that monomers are directly added to the fibrils end, without the formation of intermediate building blocks. The kinetics of fibril formation followed a sigmoidal curve, indicating nucleation dependent fibril formation. A seeding experiment with an already equilibrated sample showed accelerated kinetics, confirming catalytic activity on the fibril surface of preformed fibrils. For the titratable sidechains of H16 and H18 and the backbone proton of S21, *pK*_a_ values were determined through pH titration. H16 showed a *pK*_a_ of 4.4, which is exceptionally low for an imidazole side chain. We found that H16 in every second monomer within both strands is buried^53^, which has been shown to contribute to variability of histidine *pK*_a_ values^54^. H18 sidechain chemical shifts (and the nearby S21) showed a *pK*_a_ for the H18 sidechain of 6.4, which is very close to the pH where self-assembly becomes favorable. Thus, deprotonation of both histidine side chains parallels the pH response of fibril formation for PPI42. Through the Q14K mutation the coordination of H18 in PPI42 changed, further supporting a crucial role of H18 in the fibril formation. The reversibility of the fibril formation and the native-like protein structure within the fibril implied that only minor changes in the protein trigger its polymerization, consistent with recent reports that protein wildtypes are evolved on the edge of supramolecular self-assembly, which can be induced by few mutations^55^.

In order to investigate whether the fibrils of PPI42 could be compared to amyloid fibrils and benefit from their extensive characterization, we characterized the fibrils with a series of methods widely used for amyloid characterization^6,47,56–58^. CD spectroscopy was applied to investigate whether PPI42 undergoes structural changes upon fibril formation. We indeed observed a change in the spectrum, however, compared to amyloid fibrils formed by AMPs^21–23^, the general structural integrity of PPI42 remained intact. This is in agreement with the expected high structural identity of plectasin and the native-like structure of PPI42 in the protein fibril. Contrary to cross-α amyloid-like fibrils^36,37^ and other fibril structures formed by AMPs^20–25^ the fibrils of PPI42 proved to be negative for the fluorescence detection with ThT.

The fibril structure of plectasin variant PPI42 characterized in this study is the first of its kind, containing α-helix and β-sheet structures of the native-like protein structure. It lays the foundation for structural characterization of fibrils formed by anti-microbial peptides. Notably, the antimicrobial activity does not seem to be negatively affected by fibrillation making a slow release of bioactive monomer possible. The pH-dependent reversibility of this fibril formation and apparent anti-microbial activity of supramolecular plectasin may serve as a basis for development of novel types of anti-microbial drugs.

## ACKNOWLEDGEMENTS

This work was funded by European Union’s Horizon 2020 research and innovation program (grant agreement no. 675074).

We thank Birgitte Andersen and Ida Ahlmann Ellingsgaard from Novozymes A/S for their input on previous plectasin studies, supply of purified material, and their help on the activity measurements of plectasin.

We thank Rahmi K. Elfa for her work on the crystallization of the plectasin wildtype.

We acknowledge the provision of in-house experimental time from the CM01 facility at the ESRF. This work used the EM facilities at the Grenoble Instruct-ERIC Center (ISBG; UMS 3518 CNRS CEA-UGA-EMBL) with support from the French Infrastructure for Integrated Structural Biology (FRISBI; ANR-10-INSB-05-02) and GRAL, a project of the University Grenoble Alpes graduate school (Ecoles Universitaires de Recherche) CBH-EUR-GS (ANR-17-EURE-0003) within the Grenoble Partnership for Structural Biology. The IBS Electron Microscope facility is supported by the Auvergne Rhüne-Alpes Region, the Fonds Feder, the Fondation pour la Recherche Médicale and GIS-IBiSA.

We acknowledge MAX IV Laboratory for time on BioMAX under Proposal [20190334]. Research conducted at MAX IV, a Swedish national user facility, is supported by the Swedish Research council under contract 2018-07152, the Swedish Governmental Agency for Innovation Systems under contract 2018-04969, and Formas under contract 2019-02496

NMR spectra were recorded using the 800 MHz spectrometer at the NMR Center DTU, supported by the Villum Foundation.

## AUTHOR CONTRIBUTIONS

C.P., A.N. and P.H. designed the study. C.P. conducted, analyzed and interpreted DLS, CD, fiber diffraction and ThT experiments and RDA assay. G.E. processed cryo-EM data, and built atomic models, E.K. conducted cryo-EM experiments. G.S. and C.M.-D. assisted in cryo-EM experiments, model building and structural analysis. S.M. conducted, analyzed and interpreted NMR experiments. G.Z. conducted, analyzed and interpreted AFM experiments. P.H. conducted, analyzed and interpreted crystallization experiments. C.P. wrote the manuscript with support from A.N. and P.H. P.H., A.N., W.S. and G.H.J.P. supervised the study. All authors corrected and approved the final manuscript.

## MATERIAL AND METHODS

### Sample Preparation

If not stated otherwise, all plectasin samples were dialyzed into the desired condition using Slide-A-Lyzer™ 2000 MWCO dialysis cassettes (Thermo Fisher) with a buffer exchange after 2 and 4 hours and then continued overnight, ensuring a dilution of at least 200 times in each step. The concentration of plectasin wildtype stock solution was *c* = 39 mg/mL (8.9 mM) and of PPI42 *c* = 37 mg/mL (8.4 mM). If not gelled, the protein concentration after dialysis was measured using a NanoDrop™ 8000 Spectrophotometer. The pH reported was measured on the dialysis buffer if not stated otherwise.

### Atomic Force Microscopy

Atomic force microscopy was performed on PPI42 dialyzed in 10 mM sodium acetate pH 5, 10 mM citrate pH 5, H_2_O pH 5, 10 mM histidine pH 6.5. All samples formed a gel. A piece of gel was gently transferred to silicon wafer and dried at ambient conditions. All samples were washed three times with MilliQ water and dried again. A Multimode 8 AFM (Bruker Nano) with PeakForce Quantitative Nanomechanical Mapping (QNM) mode was used for imaging with simultaneous mapping of topography and elasticity. TAP150A probes (Budget Sensors) with nominal spring constant of 5 N/m was used. For determining elasticity of the samples, the following calibration procedure of the probe was used: 1) deflection sensitivity calibration on sapphire; 2) spring constant calibration using thermal tuning; 3) tip radius calibration using polystyrene test sample (Bruker QNM sample kit, PS film, Nominal elastic modulus 2.7 GPa); 4) calculation of elastic modulus using DMT model. The same probe was used for all measurements.

### Circular Dichroism

Plectasin was dialyzed into 10 mM sodium acetate pH 3.5 and pH 5.5 and filtered if possible (0.02 μm). A sample of 30 μM (0.132 mg/mL) was prepared by dilution with the filtered dialysis buffer. The CD spectrum was measured with a JASCO J-1000 spectrometer from 190-260 nm with 1 mm optical path. 5 spectra were accumulated for each measurement. Ellipticity θ (mdeg.) was converted to molar ellipticity [θ]:

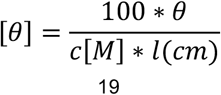

*c* is the protein concentration and *l* is the pathlength.

### Cryo-EM grid’s preparation

PPI42 was diluted with 10 mM acetate pH 5.5 to a final peptide concentration of *c*=10 mg/mL and dialyzed into 10 mM acetate pH 5.5 as described above. 3.5 μL of sample were applied to glow-discharged 2/2 Quantifoil holey carbon grids (Quantifoil Micro Tools GmbH, Germany) and plunged frozen in liquid ethane with a Vitrobot Mark IV (Thermo Fisher Scientific) (6 s blot time, blot force 0, drain time of 0.5 s). The sample was observed at the beamline CM01 of the ESRF (Grenoble, France)^59^ with a Titan Krios G3 (Thermo Fischer Scientific) at 300 kV equipped with an energy filter (Bioquantum LS/967, Gatan Inc, USA) (slit width of 20 eV). 8443 images were recorded automatically on a K2 summit direct detector (Gatan Inc., USA) in counting mode with EPU (Thermo Fisher Scientific). Movies were acquired for a total exposure of 4 s and 100 ms per frame resulting in 40 frame movies with a total dose of 46.8 e^−^/Å^2^. The magnification was 165,000x (0.827 Å/pixel at the camera level). The defocus of the images varies between −0.5 and −1.5 μm.

### Cryo-EM image analysis of the superstructure

Image processing was performed in RELION 3.1^60^ Movies were drift-corrected using MotionCor2^61^. CTF estimation of the micrographs was performed using GCTF^62^. Empty field of view and crystalline ice micrographs and micrographs with an estimated resolution lower than 4.5 Å (CTF estimation step) were removed, resulting in a set of 7241 micrographs. An initial set of particles obtained by manual picking and initial 2D class averages were used to pick automatically all micrographs using a rather conservative threshold. Improved 2D class averages for the superstructure clearly displayed an axial 2-fold symmetry. An axial rise z = 24.8 Å was measured on their Fourier transforms. An initial 3D reference was obtained by using a previously described method^63,64^, which searches iteratively for the best match between 2D projections of a 3D reconstruction computed with some given helical parameters and an ab initio 2D class average. This method identified clearly the axial rise z = 24.9 Å but failed to determine reliable azimuthal angles phi. A 3D reconstruction with z=25.1 Å and phi=16.1° was arbitrarily chosen as an initial model for 3D refinement. The latter was further low pass filtered to 40 Å to avoid any model bias. A first 3D refinement with two-fold symmetry resulted in a 3.35 Å resolution map. After two iterations of a 3D classification (without alignment) followed by a 3D refinement, a 3 Å map was calculated. The plectasin monomer was fitted unambiguously. Using the known axial rise (z = 25.1 Å), an initial azimuthal angle phi = 15.6° could be determined from the position of two successive plectasin monomers docked in the map. A second automatic picking was performed with a lower threshold to generate more particles (2 313 914 in total). The 2D classification showed a vast majority of superstructure (1 750 332 particles) as well as a small population of single fibril (99 784 particles – see below). A first 3D refinement with 2-fold and helical symmetry followed by a 3D classification and particle polishing resulted in a 2.5 Å resolution map calculated from 764 822 particles. CTF refinement in RELION was repeated two times which improved the resolution of the reconstruction to 2.05 Å resolution. A final CTF refinement (beam tilt, trefoil and 4^th^ order aberrations only) was performed with 10 000 particles per optic group to account from the variations in coma alignment during the length of the data collection. After the final 3D refinement, the final 3D map for the superstructure was calculated (final helical parameters z=25.10 Å and phi=15.75°) and a resolution of 1.97 Å was determined by Fourier Shell Correlation (FSC) at 0.143. The asymmetric unit was composed of seven monomers arranged in a near helical way with an average axial rise of 3.75 ± 0.15 Å and an average twist of 156.48 ± 2.25° between them.

### Cryo-EM image analysis of the single fibril

From the 2D classification of the superstructure, a small proportion of single fibril was identified. A separate round of 2D classification allowed the isolation of a homogenous group of single fibrils (66 272 particles). An axial rise z = 3.78 Å was measured from the fourier transform. The 3D reconstruction of the single fibril was obtained after a 3D refinement using as a starting model a cylinder of 70 Å in diameter and with initial helical parameters z = 3.75 Å and phi = 156.5°. This resulted in a 3D map at the resolution of 3.35 Å as determined by Fourier Shell Correlation (FSC) at 0.143. The final helical parameters were determined z = 3.76 Å and phi = 156.5°.

### Cryo-EM model refinement

The crystal structure of the plectasin wildtype was first rigid-body fitted inside the cryo- EM density maps in CHIMERA^65^. The atomic coordinates were then adjusted manually in COOT^66^ and refined in the cryo-EM map with ROSETTA^67^ and PHENIX^68^. The refined atomic models were visually checked and validated with MOLPROBITY^69^. The cryo-EM related figures were prepared with CHIMERA and CHIMERAX^70^. The data collection and the model statistics are summarized in Supplementary Table S3.

### Dynamic Light Scattering

Plectasin samples were dialyzed in 10 mM acetate pH 4.5 as described above, filtered (0.02 μm) and the concentration was measured. A protein stock solution of 20 mg/mL (4.5 mM) was obtained by dilution with the filtered dialysis buffer (0.02 μm). The respective formulations were obtained by a 10 times dilution. The measurement was performed with a DynaPro® Plate Reader™ II (Wyatt Technology) using Aurora 384 LV/EB plates (Brookes Life Science Systems). All measurements were performed at 25°C with 5 s acquisition time and 20 acquisitions per well. All formulations were measured in triplicates. The analysis was performed DYNAMICS version 7.8.1.3 and final graphs were made with Origin® 2019(OriginLabs).

### Fiber Diffraction

Plectasin was dialyzed into 10 mM acetate pH 5.5. The gelled sample was dried for 48 hours and was analyzed using a Supernova CCD diffractometer from Agilent.

### Negative Staining

Plectasin was dialyzed in 10 mM acetate pH 5.5. The samples were absorbed to the clean side of a carbon film on a carbon–mica interface and stained with 2% sodium silico-tungstate (pH 7.4). The carbon was transferred to a 400-mesh copper grid. Images were taken under low dose conditions (<30 e^−^/Å^2^) with defocus values between −1.2 and −2.5 μm on a Tecnai 12 LaB6 electron microscope at 120 kV accelerating voltage using CCD Camera Gatan Orius 1000.

### NMR measurements

Samples of 500 μl volume protein solution were prepared for NMR spectroscopy. The samples contained protein at 250 μM concentration (unless indicated otherwise) in 50 mM phosphate or 50 mM acetate buffer containing 10% v/v D_2_O as the lock substance. All NMR spectra were recorded on an 800 MHz Bruker Avance III NMR spectrometer equipped with an 18.7 T Oxford Magnet and a Bruker TCI CryoProbe at 298K. Protein aggregation over time was followed *in situ* by a sequence of one-dimensional ^1^H NMR experiments employing excitation sculpting as the water suppression scheme. The series of one-dimensional ^1^H NMR was implemented as a pseudo-2D experiment, which sampled 16384 complex data points during an acquisition time of 1.27 seconds and accumulated 128 transients with an interscan relaxation delay of 1 second per time point.

2D ^1^H-^1^H TOCSY NMR spectra were acquired sampling the FID for 117 and 14.5 milliseconds by acquiring 1024×128 complex data points in the direct and indirect dimension, respectively. For these ^1^H-^1^H TOCSY NMR experiments, 16 transients were acquired with an inter-scan relaxation delay of 1 second. All NMR spectra were processed with ample zero filling in all spectral dimensions in Bruker Topspin 3.5 pl7 software and integrated in the same software. Data were plotted in proFit 7 (Quantum Soft, Switzerland). Titration curves were fitted to the Henderson-Hasselbalch equation in proFit 7 as

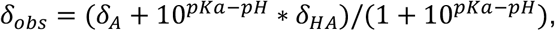

where δ_obs_ is the measured chemical shift, δ_A_ is the chemical shift of the deprotonated form and δ_HA_ is the chemical shift of the protonated form. The pH of the sample was determined after the measurement.

### Radial Diffusion Assay

Preparation of target strain, *Staphylococcus carnosus*: Stock of target strain was incubated at 37°C overnight in anaerobic conditions (AnaeroGen 2.5L, Thermo Scientific) on to LB agar plate then resuspended in 0.9% NaCl, 20% glycerol to a McFarland turbidity standard of 1 (OD_600_ approx. 0.870). Aliquots of 150 μl are frozen at −80°C. The colony forming unit per mL (CFU/mL) was determined. The RDA media (5.5 g Mueller Hinton II Broth, Beckton Dickinson; 7.5 g Agarose High resolution, Sigma Aldrich, 500 ml MilliQ water) was autoclaved (121°C; 15 minutes) and cooled down to approx. 42°C. *S. carnosus* was added 30 ml medium to a final CFU/mL of 5×10e^5^. Plates were prepared using Omnitray with NUNC-TSP lid (both Thermo Scientific) and kept at 4°C for least 20 minutes to solidify. The lid was discarded, and the wells were loaded with 10 μl sample. The RDA plate was incubated at 37°C overnight. The plate was colored with 1.5 mM Thiazolyl Blue Tetrazolium Bromide (MTT) (Sigma Aldrich).

### Thioflavin T Fluorescence Measurements

Plectasin was dialyzed in 10 mM acetate pH 5.5. A final concentration of 2 mg/mL was obtained by dilution with the dialysis buffer and ThT was added to a final concentration of 20 μM. The fluorescence was monitored after each cycle at 490±10 nm with excitation at 440±10 nm with a gain of 1000. Insulin was fibrillated after the protocol of Frankjær et al^46^. Additionally, the fluorescence of a sample containing insulin fibrils at a peptide concentration of 2 mg/mL was compared to freshly dialyzed plectasin in 10 mM sodium acetate buffer pH 3.5 and pH 5.5 and a sample diluted into 10 mM sodium acetate buffer pH 5.5 and 10 mM phosphate buffer pH 6.5, equilibrated for 12 hours was compared (end-point measurement).

### X-ray Crystallography

#### Crystallization

Crystallization conditions of plectasin were screened using a sparse matrix screen^71^ by Molecular Dimensions. Crystallization conditions were optimized. Crystals formed overnight.

#### Data collection and structure determination

Crystals were flash cooled directly in liquid nitrogen. X-ray data was collected at 100 K at the BioMAX beamline at MAXIV, Lund, Sweden^72^. Data reduction was performed with the autoPROC toolbox^73^ using XDS/XSCALE^74^ and Pointless^75^.

#### X-ray structure determination

The crystal structure was determined using the CCP4 program suite^76^ with molecular replacement using Molrep^77^. The previously solved crystal structure of plectasin 3E7U^43^ was used as template. Restrained positional and anisotropic B-factor refinement was performed in REFMAC5^78^. The hydrogen atoms were included in riding positions. Data collection and refinement statistics are given in Table S4.

