## Supplementary Information for "Reversible supramolecular assembly of the anti-microbial peptide plectasin into helical non-amyloid fibrils"

Table S1: Thioflavin T fluorescence end-point measurement on plectasin wildtype, PPI42 and insulin (c=2 mg/mL) in different conditions (excitation: 440±10 nm; emission: 490±10 nm).

| sample | buffer | preparation | fluorescence (490 nm) |
|---|---|---|---|
| Insulin | Gly-HCl pH 2.5 | dissolved | 13428.8 |
| wildtype<br>PPI42 | Acetate pH 3.5 | dialyzed | -36.4<br>-37.4 |
| wildtype<br>PPI42 | Acetate pH 5.5 | dialyzed | 6.4<br>-12.2 |
| wildtype<br>PPI42 | Acetate pH 5.5 | diluted | -15.2<br>-13.8 |
| wildtype<br>PPI42 | Phosphate pH 6.5 | diluted | 57.8<br>-1.8 |

Table S2: Ionic strength of buffers used in this study.

| | ionic strength (mM) | | | |
|---|---|---|---|---|
| pH | Acetate | Phosphate | Histidine | Citrate |
| 3.5 | 0.52 | | | 7.92 |
| 4.0 | 1.49 | | | 11.68 |
| 4.5 | 3.56 | | | 16.83 |
| 5.0 | 6.36 | 10.26 | 9.13 | 23.49 |
| 5.5 | 8.47 | 10.82 | 7.61 | 30.05 |
| 6.0 | 9.46 | 12.4 | 5 | 37.32 |
| 6.5 | | 16.03 | 2.4 | |
| 7.0 | | 21.54 | 0.91 | |
| 7.5 | | 26.24 | 0.3 | |
| 8.0 | | 28.64 | 0.24 | |

Table S3: Summary of cryo-EM data collection and atomic model statistics of PPI42.

| Data collection | | |
|---|---|---|
| Microscope | Krios G3 (ThermoFischer Scientific) | |
| Voltage (kV) | 300 | |
| Magnification<br>Unbinned pixel size | 165,000x<br>0.827 Å/pixel | |
| Camera | K2 Summit (Gatan Inc) | |
| Exposure time | 4s | |
| Number of frames | 40 | |
| Total dose ($e^{-}/Å^{2}$) | 46.4 | |
| **Image processing** | | |
| | Superstructure | Single fibril |
| EMDB | EMD-12775 | EMD-12776 |
| Rotational symmetry | C2 | C3 |
| Helical symmetry (axial rise/azimuthal angle) | 25.1 Å / 16.75° | 3.76 Å / 156.5° |
| Final number of Particles | 764822 | 66272 |
| Map resolution in Å (FSC 0.143) | 1.97 | 3.35 |
| **Model statistics** | | |
| PDB | 7OAE | 7OAG |
| Model resolution in Å (FSC 0.5) | 2.15 | 3.5 |
| Ramachandran favored (%) | 97.4 | 94.7 |
| Ramachandran outliers (%) | 0.0 | 0 |
| Rama Z score | 0.22 | -0.35 |
| Rotamer outliers (%) | 0.0 | 0 |
| C-beta deviations | 0 | 0 |
| Rms on bond lengths | 0.0076 | 0.0069 |
| Rms on bond angles | 0.53 | 0.67 |
| Clashscore | 0.99 | 5.52 |
| Molprobity score | 0.91 | 1.66 |

Table S4: Summary of X-ray data collection and refinement of WT plectasin.

| **Data collection** | |
|---|---|
| Space group | $P2_1$ |
| Unit cell | |
| a, b, c (Å) | 23.68, 20.09, 32.55 |
| α, β, γ (°) | α= 90, β=104.28, γ=90 |
| Total number of reflections | 59453(3272) |

| | |
|---|---|
| Number of unique reflections | 10216(1020) |
| Protein molecules in ASU | 1 |
| Resolution limits (Å) | 31.53-1.13(1.20-1.13) |
| $R_{merge}$ | 0.033(0.343) |
| $R_{pim}$ | 0.018(0.274) |
| CC(½) | 0.999(0.900) |
| Completeness (%) | 89.9(54-6) |
| Average I/σ (I) | 22.49 (2.65) |
| Wilson plot B-factor (Å$^2$) | 17.38 |
| Multiplicity | 6.0(3.6) |
| **Refinement** | |
| $R_{work}$ | 0.1459 |
| $R_{free}$ | 0.1762 |
| Number of reflections | 9708 |
| Reflections used for R-free | 507 |
| Number of non-hydrogen atoms in ASU | 320 |
| Number of water molecules | 25 |
| No non-hydrogen atoms in ligand | 7 |
| **Root mean square deviation from ideal** | |
| Bond lengths (Å) | 0.018 |
| Bond angles (°) | 2.005 |
| B-factors (Å$^2$) | 26.43 |
| Solvent content | |

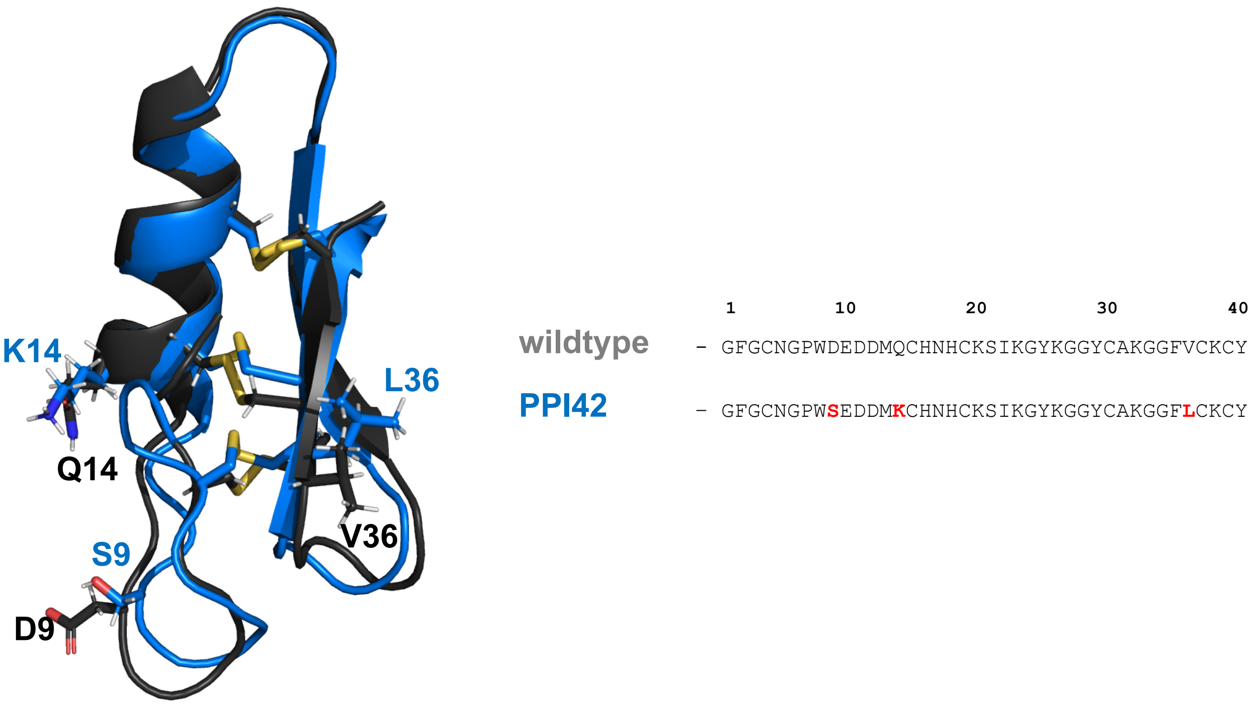


Figure S1: Structure of plectasin wildtype (black) (PBDID: 3E7U[43]) and energy-minimized structure of PPI42 (blue) with mutated amino acids and three disulfide bonds (C4-C30; C15-C37; C19-C39) shown as sticks. The sequence is shown with mutated amino acids labeled in red for PPI42. Figure was made with PyMOL[79].

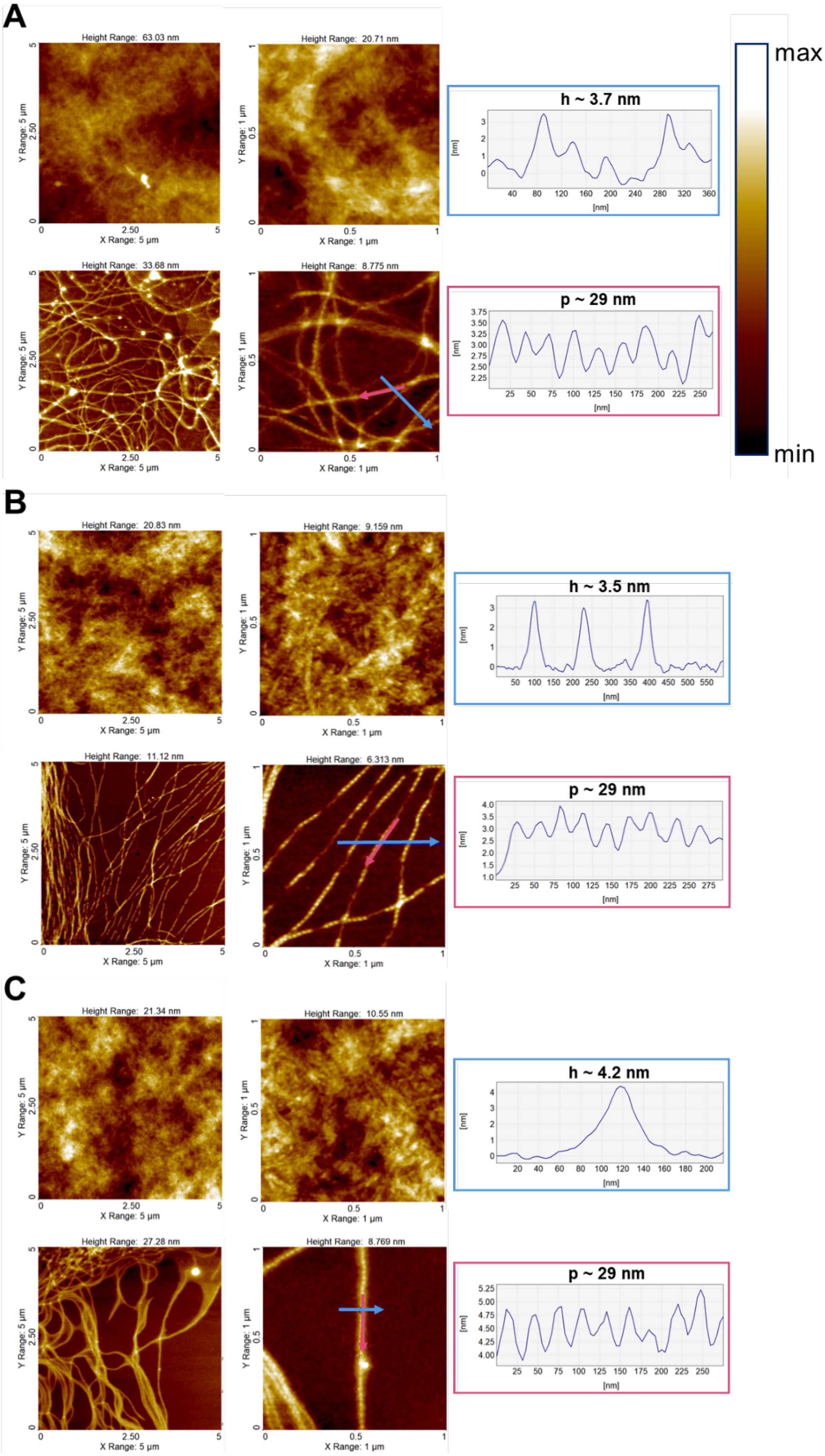


Figure S2: A: AFM measurements on PPI42 fibrils in histidine buffer pH 6.5. B: AFM measurements on PPI42 fibrils in acetate buffer pH 5. C: AFM measurements of PPI42 fibrils in $H_2O$ pH ≈ 5. The respective measured height (h) and the periodicity along the fibril (p) are shown on the right.

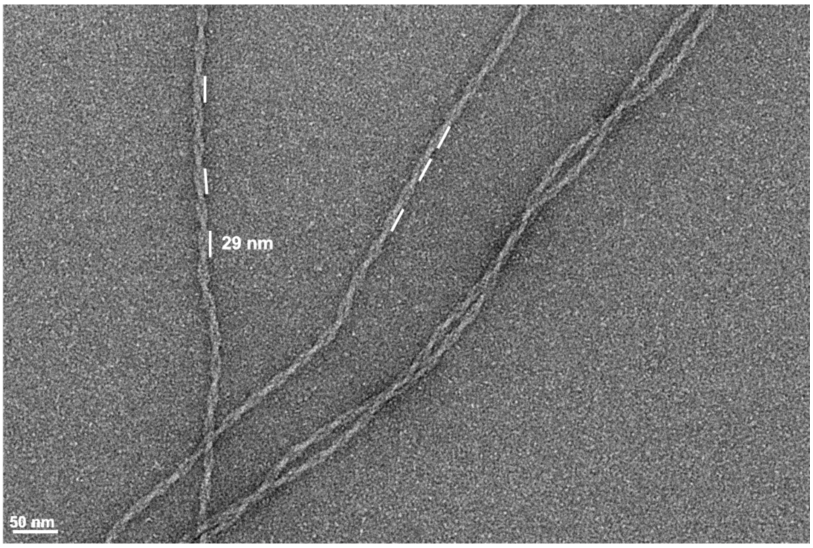

Figure S3: Exemplary measurement of periodicity along the fibril of images obtained with negative stain EM.

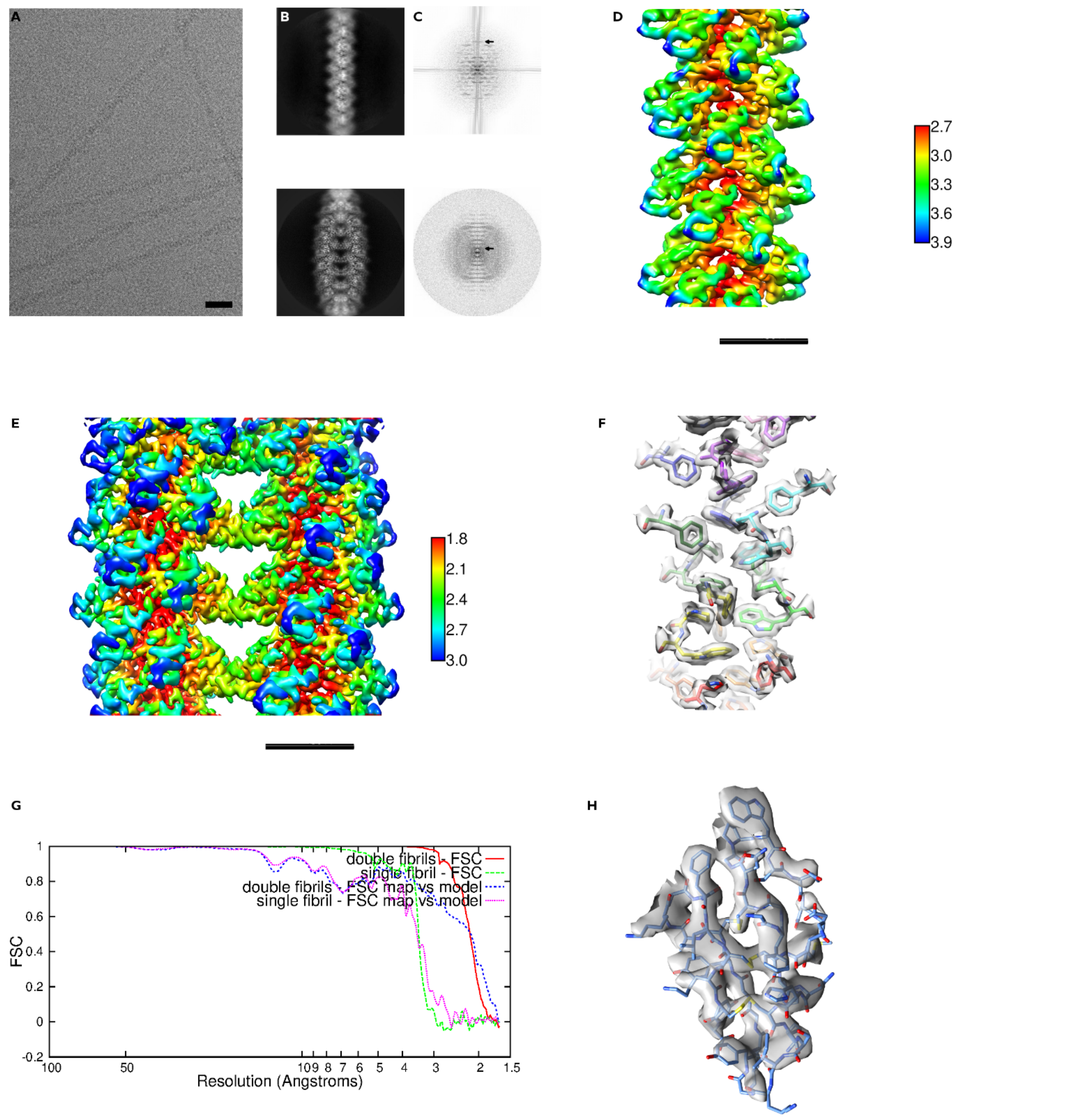


Figure S4: A: Cryo-EM field of view of PPI42 fibrils. B: Exemplary 2D class averages. Upper panel: single fibrils; lower panel: fibril superstructure. C: Power spectra of the two class averages shown in B. Arrows point to layer lines at 24.8 Å and 3.8 Å for the single fibrils and fibril superstructure, respectively. D and E: Local resolution maps for the single fibrils (D) and superstructure (E) reconstructions. Color code indicate resolution in Å. The scale bar represents 30 Å. F: Illustration of the high-resolution features of the fibril superstructure reconstruction displaying the hydrophobic core (P7, W8 and F35) of the superstructure. PPI42 monomers are colored individually. Cryo-EM Coulomb potential map is shown in gray. G: Fourier Shell Correlation (FSC) curves: gold standard FSCs between two independent 3D reconstructions of the single fibril (green dotted line) and fibril superstructure (red line) and FSC between the cryo-EM maps and the refined atomic models for both the single fibril (dotted pink line) and fibril superstructure (dotted blue line). H: Atomic model of PPI42 monomer built in the single fibril reconstruction (3.4 Å overall resolution).

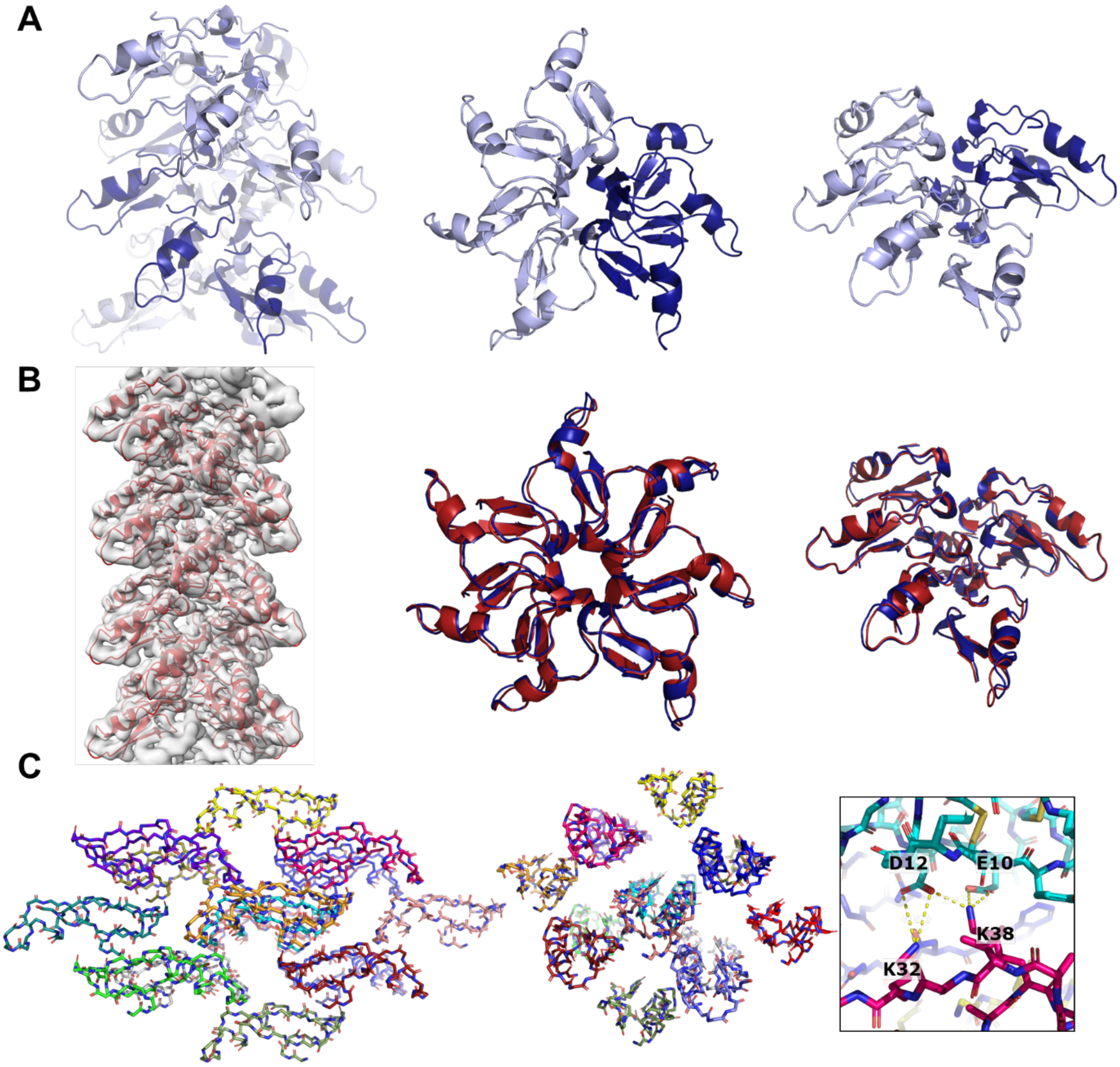


Figure S5: A: Double stranded near-helical architecture of fibril in the superstructure formed by PPI42 (left) and asymmetric unit of the fibril shown from the top and from the front (right). The two strands are colored in different shades of blue. B: Cryo-EM map of the single fibril at a resolution of 3.4 Å and build in atomic structure of PPI42 monomers. C: Crystal arrangement of the plectasin wildtype at 1.1 Å resolution from the front and from the top (left). Conserved crystal contacts of the acidic amino acid stretch D9, E10, D11, D12 are shown in detail (right).

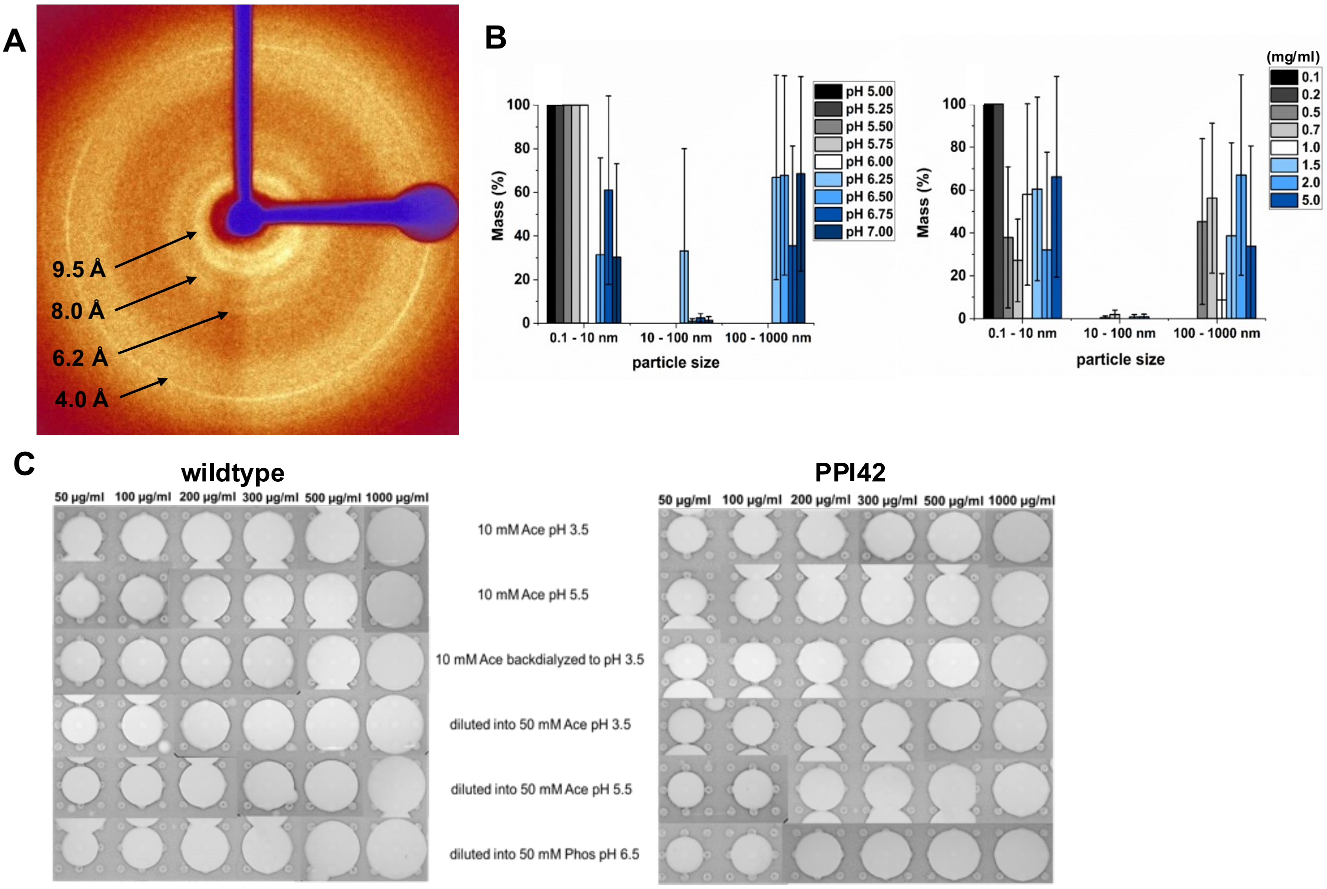


Figure S6: A: Fiber diffraction image of the plectasin fibrils of PPI42. The relevant diffraction signals and distances are highlighted. B: Dependence of PPI42 oligomerization on protein concentration determined by DLS shown as %Mass at pH 6.5. Data is mean ± S.D. for 3 replicates. C: Clearing zones to measure the anti-microbial activity of the plectasin wildtype and PPI42 as a function of peptide concentration in different conditions.

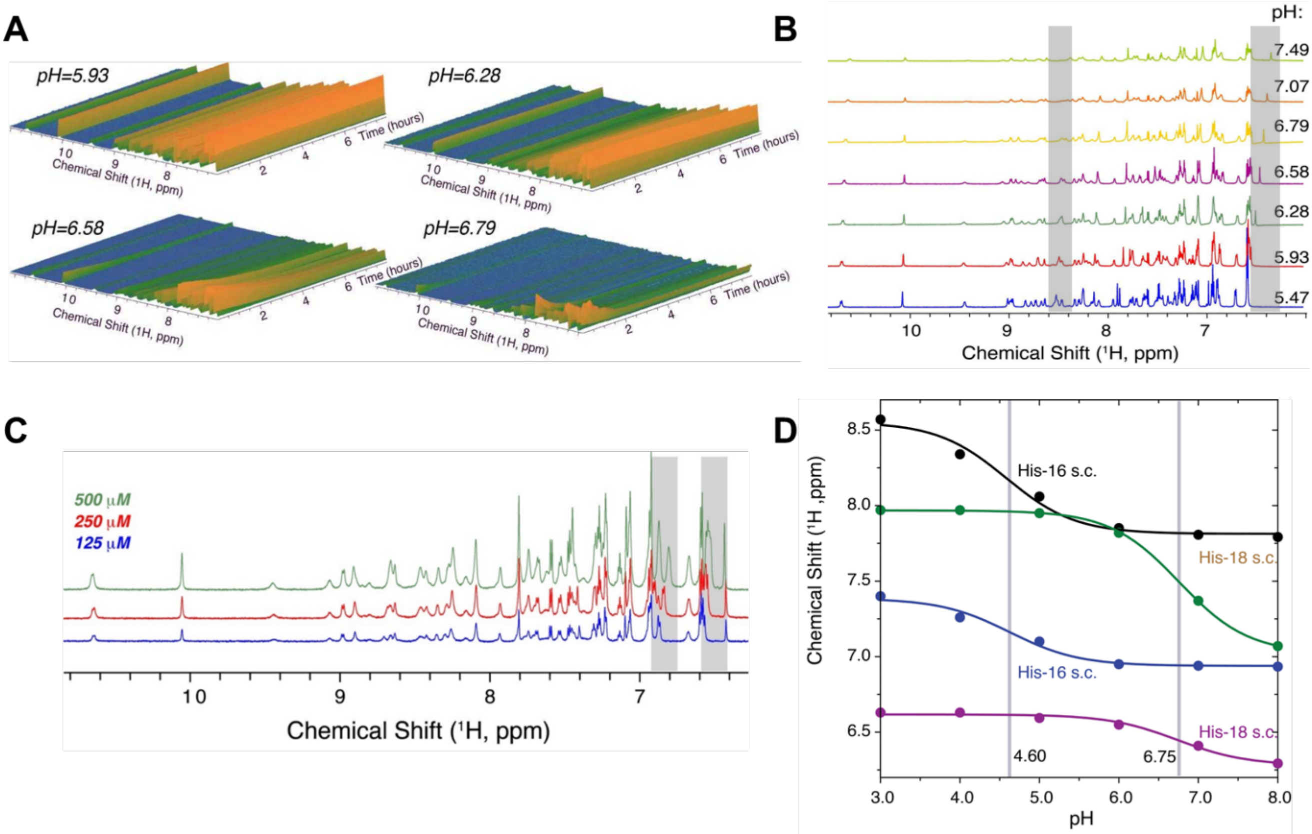


Figure S7: A: Loss of monomer signals in NMR over time for PPI42 (c=0.25 µM, 298 K) at different pH (50 mM phosphate buffer). B: $^{1}$H chemical shift of PPI42 in a series of pH from pH 5.5 to 7.5. C: $^{1}$H NMR spectra of PPI42 at pH 6.75 at different protein concentrations. The area, where a change in NMR signals is most evident is shaded in grey. D: Titration curves of wildtype plectasin H16, H18 ($H^2$ and $H^4$ in imidazole ring) and S21 $H^N$ determined by resolving the signals with 2D $^{1}$H-$^{1}$H TOCSY. The grey bars indicate the $pK_a$ of the titratable side chains. All measurements were conducted in 50 mM phosphate buffer.
